# Regenerative Index reveals declining muscle regeneration in paediatric patients with Duchenne muscular dystrophy

**DOI:** 10.64898/2026.01.05.697715

**Authors:** Johnathan K. Smid, Charis A. McPherson, Jacob G. Monast, Shanti S. S. Rayagiri, Steven A. Moore, Michael A. Rudnicki

**Affiliations:** Sprott Centre for Stem Cell Research, Regenerative Medicine Program, Ottawa Hospital Research Institute, Ottawa, Canada; Department of Pathology, Carver College of Medicine, University of Iowa, Iowa City, USA; Department of Cellular and Molecular Medicine, Faculty of Medicine, University of Ottawa, Ottawa, Canada; Satellos Bioscience, Toronto, Canada

**Keywords:** Regenerative Index (RI), Muscle regeneration, Duchenne muscular dystrophy (DMD), dystrophin, Embryonic myosin heavy chain (eMHC), Regenerating myofibers, Necrotic myofibers

## Abstract

**Background:** Duchenne muscular dystrophy (DMD) is a devastating disease manifested in skeletal muscle by repetitious myonecrosis and regeneration. Because the regenerative process is closely linked to the cumulative severity of muscle damage, which is variably distributed within and between muscle groups, accurately quantifying muscle regeneration has remained a significant challenge.

**Methods:** Myofibers are delineated by immunostaining for laminin, and subsequent image analysis employed to generate a masked outline precisely within each myofiber boundary. Morphometric parameters including minimal Feret’s diameter, cross-sectional area, and circularity were measured for each myofiber. In addition, the number of Pax7-expressing satellite cells were quantified. To evaluate regenerative activity, newly formed myofibers were identified by immunostaining for expression of embryonic myosin heavy chain (eMHC). Necrotic myofibers were enumerated by immunofluorescent detection of immunoglobulin G (IgG) infiltration. The Regenerative Index (RI) was calculated as the number of regenerating (eMHC^+^) myofibers divided by the number of necrotic (IgG^+^) myofibers. Determination of RI was performed on muscle biopsies from 10 boys with DMD and 3 non-DMD controls of similar age.

**Results:** A trend toward an increasing minimal Feret’s diameter, cross-sectional area and circularity was observed with increasing age in DMD boys, with circularity showing the strongest trend. Furthermore, compared to DMD boys 7- to 8-years old, the boys 9- to 11-years old had significantly increased myofiber circularity. Pax7-expressing cells were significantly elevated in DMD boys compared to control boys of similar ages, without any observation of age-related changes. Notably, the Regenerative Index in DMD boys exhibited a pronounced decline between 7-11 years of age, and a significant inverse correlation between RI and age was observed.

**Conclusions:** Using eMHC and IgG immunostaining to calculate RI accurately assesses regeneration despite the variation in histopathologic severity between biopsies. This methodology demonstrated a significant negative correlation between RI and age of DMD boys from 7 to 11 years of age.

## Background

Duchenne muscular dystrophy (DMD) is a severe X-linked neuromuscular disorder caused by pathogenic variants in the *dystrophin* gene (*DMD*) leading to an absence or non-functional dystrophin protein [1, 2]. Although still considered a rare disease, DMD is one of the most common forms of muscular dystrophy with a prevalence of 1:5,000 boys [3, 4]. Furthermore, it is one of the most severe forms of muscular dystrophy with a dramatic decline in abilities at a young age and a loss of independent walking in the pre-teen to early teenage years [5]. Although current standards of care and emerging gene-targeting therapies for DMD can prolong ambulation and improve survival into adulthood, the disease remains relentlessly progressive, and most continue to experience life-limiting respiratory and/or cardiac complications preventing a normal lifespan [6]. Taken together, there is an urgent need for more effective and widely accessible treatments for DMD.

Previous studies characterizing the trajectory of DMD myopathology have highlighted the lack of muscle regeneration [7, 8]. Moreover, intrinsic deficits from the lack of dystrophin in muscle stem cells has been shown to be the cause [9–13]. In *mdx* mice, the mouse model of DMD, the lack of dystrophin results in reduced polarity and the loss of asymmetric divisions [9], a key mechanism through which muscle stem cells maintain a pool of progenitor cells to facilitate regeneration [14]. This impairment results in reduced muscle regeneration, the senescence of muscle stem cells, and an accumulation of fibro-adipogenic progenitors [7, 15, 16]. A treatment to rescue this satellite cell defect and restore regeneration holds promise to be of therapeutic benefit [17].

A method to determine the efficacy of regeneration-enhancing therapeutics is currently lacking. To properly evaluate the effectiveness of such treatments, a biomarker specific to the quantification of muscle regeneration in response to degeneration would be critical. Biomarkers commonly employed in DMD trials, such as such as levels of muscle fiber dystrophin or reductions in markers of muscle fiber injury, are inadequate for evaluating interventions that enhance muscle regeneration, as these endpoints fail to capture improvements in regenerative capacity.

Evaluation of the general appearance of muscle using Haematoxylin and Eosin (H&E) staining has been the gold standard for muscle tissue assessment, as it can provide information on both degenerative and regenerative processes. Degeneration can be denoted by myonecrosis, the inflammatory response, and endomysial fibrosis, while manifestations of regeneration include basophilic sarcoplasm, increased variability in myofiber size, and internally-placed nuclei [18]. Nevertheless, H&E has several caveats when used alone for evaluating muscle degeneration and regeneration. Interpretation can be subjective and dependent on observer experience, which can introduce variability in scoring both myonecrosis and regeneration [18]. Furthermore, the multifocal nature of muscle damage, and variability within and between muscles compounds the challenge. Thus, a robust, objective method for quantifying both muscle regeneration and degeneration is essential to minimize subjectivity in scoring.

Embryonic myosin heavy chain (eMHC) has long been used to identify newly forming myofibers [19]. Moreover, quantification of the percentage of regenerating myofibers reveals a negative correlation with functional motor score in DMD and BMD patients [20]. Evaluation of regeneration alone does not provide a complete representation as it comes in response to the myonecrosis occurring in these patients.

Myofiber necrosis can be quantified by the immunodetection of Immunoglobulin G (IgG) infiltration into myofibers due to loss of membrane integrity [21]. However, extensive variability in muscle fiber loss resulting from the variable myonecrosis and regeneration within and between muscles limits the utility of any determination of the rate of regeneration. To address this issue, we developed and validated a measure of muscle regenerative capacity, which we term the Regenerative Index (RI). Using this approach, regeneration is normalized to myonecrosis effectively reducing this variability. The RI is the ratio of newly formed (eMHC^+^) myofibers over necrotic (IgG^+^) myofibers, thus normalizing the variation found between biopsies. Accordingly, the RI integrates both regenerative and degenerative states to assess the net regenerative potential of muscle. We propose that determination of RI will allow the monitoring of changes in the ability of DMD muscle to regenerate, providing a powerful tool to assess treatments enhancing muscle regeneration.

## Methods

To develop a method to quantify a regenerative index in DMD and non-DMD, no histopathologic diagnostic abnormality controls (CTRL), muscle biopsy cryosections were stained and quantified using the methods outlined below. All muscle samples from biopsies performed at ages of 7 to 11 years old were obtained from the Repository at the University of Iowa Wellstone Muscular Dystrophy Specialized Research Center (IRB ID#200510769; original approval on 02/16/2006; most recent continuing review approval on 11/11/2025; see Table 1). Samples were received as frozen 10 µm sections on glass slides. All sections remained frozen until the time of immunostaining.

**Table 1:** List of muscle samples with age, muscle type and *DMD* variant, if known.

| Sample ID | Age | Muscle | *DMD* variant |
| --- | --- | --- | --- |
| DMD18 | 7 | Quadriceps | not known |
| DMD29a | 7 | Quadriceps | c.4237delA, p.Ile1413Serfs*5 |
| DMD33a | 8 | Quadriceps | 29kb deletion of intron 44 and all of exon 45 |
| DMD58a | 8 | Quadriceps | exon 19 out of frame deletion that causes frameshift p.Ala765Argfs*15 |
| DMD5a | 8 | Quadriceps | not known |
| DMD36 | 9 | Quadriceps | not known |
| DMD9 | 9 | Anterior tibialis | c.457C>T, p.Gln153* |
| DMD28 | 10 | Unknown | not known |
| DMD44b | 10 | Quadriceps | ex 38-43 out of frame deletion that causes a frameshift p.Ser1777Ilefs*2 |
| DMD32 | 11 | Anterior tibialis | c.457C>T, p.Gln153* |
| CTRL12 | 8 | Quadriceps | none |
| CTRL77a | 10 | Quadriceps | none |
| CTRL76 | 11 | Unknown | none |

DMD- Duchenne muscular dystrophy; CTRL- Control (non-DMD)

### Dystrophin Immunofluorescence

To determine the presence or absence of dystrophin msIgG2b Dystrophin 4C7 (Santa Cruz, Cat#sc-33697) was prepared at a 1:100 dilution. This antibody binds to an epitope at the n-terminus of dystrophin that is coded by exon 1 of the *DMD* gene. Slides were incubated with primary antibodies for 1hr at RT followed by three washes with phosphate-buffered saline (PBS). The secondary antibody Goat Anti-mouse IgG2b Alexa Fluor 488 (Catalog# 21141) was prepared in PBS at a concentration of 1:1000. Secondary antibodies were incubated at RT for 30 minutes. After incubation slides were washed five times with PBS before adding 4′,6-diamidino-2-phenylindole (DAPI) for 5 minutes. After coverslipping slides with PermaFluor aqueous mounting medium (Fisher scientific, cat# TA030FM), each complete muscle biopsy section was scanned on a Zeiss Axio Observer Z1/7 microscope. In parallel, non-DMD sections were stained as a positive control (Supplemental Figure 1).

### Determination of Regenerative Index by Immunofluorescence

To identify IgG^+^ and eMHC^+^ myofibers Mouse IgG2a anti-human IgG mAb (Abcam, Cat# ab200699) and Mouse IgG1 anti-eMHC mAb (Santa Cruz, Cat# sc-53091) were prepared in PBS at a dilution of 1:200. In addition, to identify myofiber membranes Rabbit Anti-Laminin (Sigma, Cat# L9393; binds to laminin 111) was also added at a dilution of 1:1000. Slides were incubated with primary antibodies for 1hr at room temperature (RT) followed by three washes with PBS. The secondary antibodies Goat Anti-Mouse IgG2a Alexa Fluor 488 (Invitrogen, Cat# A21131), Goat Anti-Mouse IgG1 Alexa Fluor 546 (Invitrogen, Cat# A21123) and Goat Anti-Rabbit Alexa Fluor 647 (Invitrogen, Cat# A21244) were prepared in PBS at a concentration of 1:2000. Secondary antibodies were incubated at RT for 30 minutes. After incubation slides are washed five times with PBS before adding DAPI for 5mins. After mounting slides the complete muscle biopsy was scanned on a Zeiss Axio Observer Z1/7 microscope. In parallel, sections were stained with secondary antibodies only to validate the specificity of staining (Supplemental Figure 2).

### Morphometric Analysis of Myofibers

Zen Image analysis software (Zeiss Zen version 3.7) was used to quantify myofiber dimensions. Briefly, the basal lamina border of each myofiber was delineated by fluorescent laminin immunostaining (Figure 1a,b) and image analysis software provided a trace within the laminin staining (Figure 1c). Zen image analysis software provided the cross-sectional area (CSA), minimal fiber ferret (MFF) and circularity of each myofiber. The quantification is semi-automated, with the script automatically performing all traces according to operator specifications, followed by minor manual edits to split myofibers that the script has merged or add in parts of the myofiber which were mis-selected. Further analysis of data exported from the Zeiss image analysis software was carried out with Graphpad Prism (version 10.4.1) and Microsoft Excel (version 16.89).

**Figure 1.**
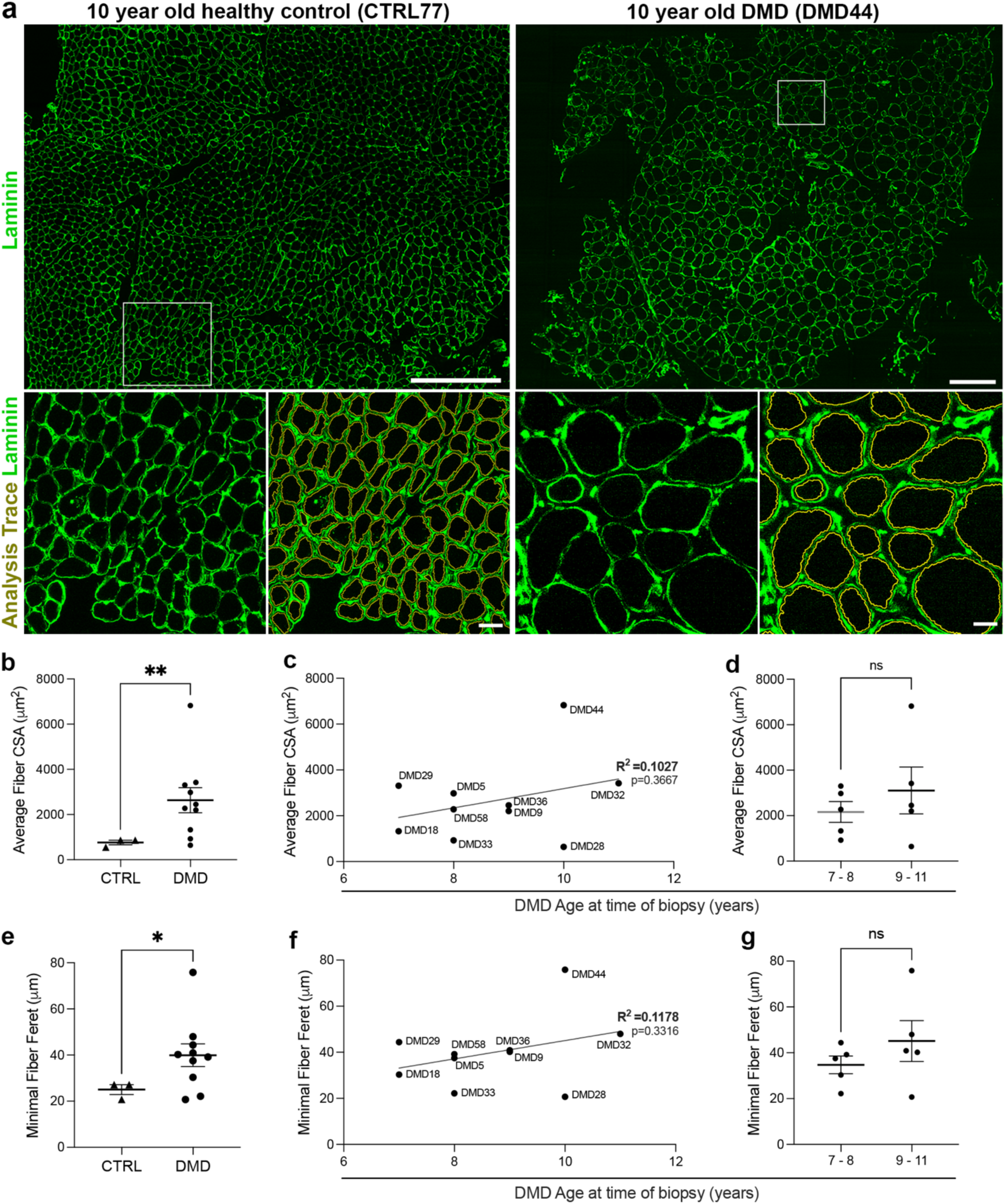
Increased fiber size in DMD muscle samples relative to non-DMD controls. (**a**) Representative laminin 111 fluorescent immunostaining (green) of a 10-year-old healthy control (CTRL77) and an aged-matched DMD patient (DMD44), with 500 µm insets showing the corresponding analysis traces to illustrate automated segmentation of individual fibers. (**b**) Quantification of average fiber cross-sectional area (CSA) shows significantly larger fibers in DMD compared with non-DMD controls (**p < 0.01, unpaired t test). (**c**) Average fiber CSA plotted against age at biopsy in DMD patients reveals a weak, non-significant positive correlation (R^2^ = 0.1027, p = 0.3667; simple linear regression). (**d**) Similarly, comparison of younger (7–8 years) and older (9–11 years) DMD patients shows no significant difference in average fiber CSA (ns, unpaired t test). (**e**) DMD muscle samples exhibit a significant increase in minimal fiber Feret diameter (MFF) compared to non-DMD controls (*p < 0.05, unpaired t test). (**f**) Correlation between MFF and age remains weak (R^2^ = 0.1178) and has a non-significant correlation (p = 0.3316; simple linear regression). (**g**) Likewise, there is no significant difference in MFF observed when comparing younger (7–8 years) and older (9–11 years) DMD patients (ns, unpaired t test).

### Pax7 Immunofluorescence and Quantification

To identify PAX7-expressing satellite cells mouse IgG1 anti-Pax7 hybridroma (DSHB) was thawed at 37C and vortexed for 20 seconds. Rabbit anti-Laminin (Sigma, cat#L9393) was added to the Pax7 hybridoma for a final concentration of 1:1,000. Slides were incubated with primary antibodies for 1hr at room temperature (RT) followed by four washes with PBS. The secondary antibodies anti-rabbit A488 (Invitrogen, cat# A11008) and anti-mouse IgG1 A546 (Invitrogen, cat# A21123) were prepared in PBS at a concentration of 1:1000. Secondary antibodies were incubated at RT for 30 minutes. After incubation slides were washed five times with PBS, stained for DAPI, and mounted with coverslips as described above. The complete muscle biopsy section was scanned on a Zeiss Axio Observer Z1/7 microscope. Quantification was performed with Zeiss image analysis software by first identifying the nucleus with DAPI and then setting a threshold to determine if the nuclei is Pax7^+^. Quantification was performed in a semi-automated manner, with each section visually inspected to identify false positives in regions of high background.

### Regenerative Index (RI) measurement

Immunofluorescent staining for laminin was combined with staining for Human IgG and eMHC. After an automated trace of the laminin border performed with Zen image analysis software the fluorescent intensity within the myofiber boundery was measured with the same software to identify either IgG+ or eMHC+ myofibers. A representative image of a quadricep muscle is shown stained with IgG and eMHC (Fig 4a). The total number of myofibers (class 1) and those expressing eMHC (class 2) and IgG (class 3) were enumerated using a script developed with the Zen software. This allowed for the calculation of the individual percentage of both the eMHC+ and IgG+ over total number of myofibers. Moreover, this allows for the Regenerative Index (RI) to be calculated as the total number of eMHC immunostained myofibers divided by the total number of IgG immunostained myofibers.

### Statistical Analysis

Statistical comparisons between two groups were performed using an unpaired Welch’s two-tailed t-test to account for potential differences in variance. For correlative studies, simple linear regression was employed to calculate the coefficient of determination (R^2^) and to assess whether the regression slope differed significantly from zero. All statistical analysis was carried out with Graphpad Prism (version 10.4.1).

## Results

### Histological analysis of disease progression in DMD

We conducted our analysis on DMD patients from 7- to 11-years of age because these patients are the focus of many clinical trials [22] as they typically are initially ambulatory and lose ambulation within four years [5, 23]. Representative H&E photomicrographs from each DMD biopsy and from one non-DMD control are shown in Supplemental Figure 3. We examined the standard markers for disease progression including cross-sectional area (CSA), mean fiber Feret (MFF), and circularity of myofibers [18, 24–26]. Moreover, we examined the abundance of satellite cells by immunofluorescent detection of Pax7 to gain insight into the regenerative process.

DMD muscle samples displayed a significantly elevated average myofiber cross-sectional area (CSA) compared to age-matched healthy muscle biopsies (Figure 1a, b). Although myofiber CSA demonstrated a trend toward increased values with age, variability across DMD samples yielded a weak correlation (low R^2^ value) with age, and the regression slope did not significantly differ from zero (Figure 1c). In addition, comparing the average of all five 7–8-year-old DMD samples to the average of five samples from 8-11 years of age did not show a statistically significant difference (Figure 1d).

The minimal fiber Feret (MFF) or the minimal Feret’s diameter is the smallest distance between two parallel tangents to the myofiber cross-section and has been used as a robust measure of myofiber size as the error from non-perpendicular sectioning is reduced [27]. Similarly to CSA, the MFF of DMD samples showed a significant increase when compared to healthy controls (Figure 1e). However, despite a trend toward increased MFF with age, substantial variability resulted in a weak age-MFF correlation (low R^2^) with a non-significant regression slope (Figure 1f). Moreover, no statistically significant difference in MFF was observed between DMD samples from 7–8-year-old versus 9-11-year-old patients (Figure 1g).

Examination of circularity revealed that there was a statistically significant shift in distribution of circularity in older versus younger boys with a stronger correlation than MFF or CSA (Figure 2). Notably, the mean circularity was higher in muscle biopsies from 9–11-year-olds compared with 7–8-year-old DMD boys (Figure 2b,c). Moreover, increased circularity was observed in areas where myofibers were less compact (Figure 2a, bottom insets) when compared to more tightly packed myofibers (Figure 2a, top insets). However, the high variability observed in younger boys resulted in a weak correlation with a very low R^2^ value and non-significant regression analysis (Figure 2d).

**Figure 2.**
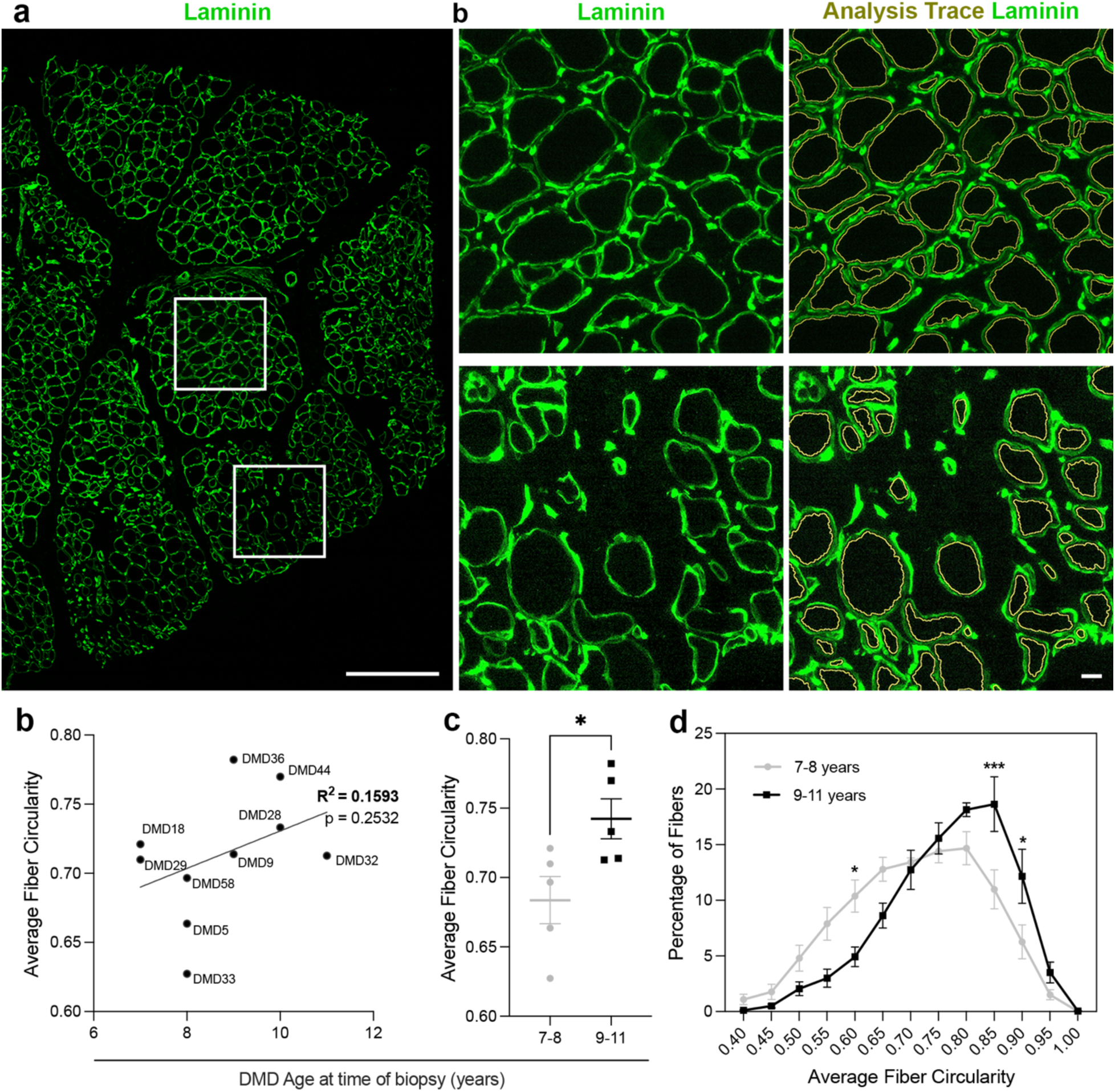
Myofiber circularity is significantly increased with age in DMD muscle. (**a**) Representative laminin-immunostained DMD muscle section showing heterogeneous fiber shape with 500 µm insets including the image analysis trace in yellow. (**b**) Distribution of average fiber circularity in younger (7–8 years, grey) and older (9–11 years, black) DMD patients, showing a rightward shift toward more circular fibers in the older group. (**c**) Mean fiber circularity is significantly increased in older compared with younger DMD patients (unpaired t test, *p < 0.05). (**d**) Average fiber circularity shows a weak, non-significant positive correlation with age at biopsy in DMD patients (simple linear regression, R^2^ = 0.1593, p = 0.2532).

PAX7 staining was combined with laminin to enumerate satellite cells in both DMD and non-DMD control (CTRL) muscle samples from boys 7-11 years of age (Figure 3a). Quantifying the number of PAX7-expressing cells and the number of myofibers revealed a significant increase in the number of PAX7-expressing satellite cells in DMD samples relative non-DMD controls (Figure 3b). Moreover, there was no decrease in satellite cell number as the boys aged and in fact there was a trend of an increase (Figure 3c), without any significant difference when comparing 7–8 years with 9-11 years (Figure 3d).

**Figure 3.**
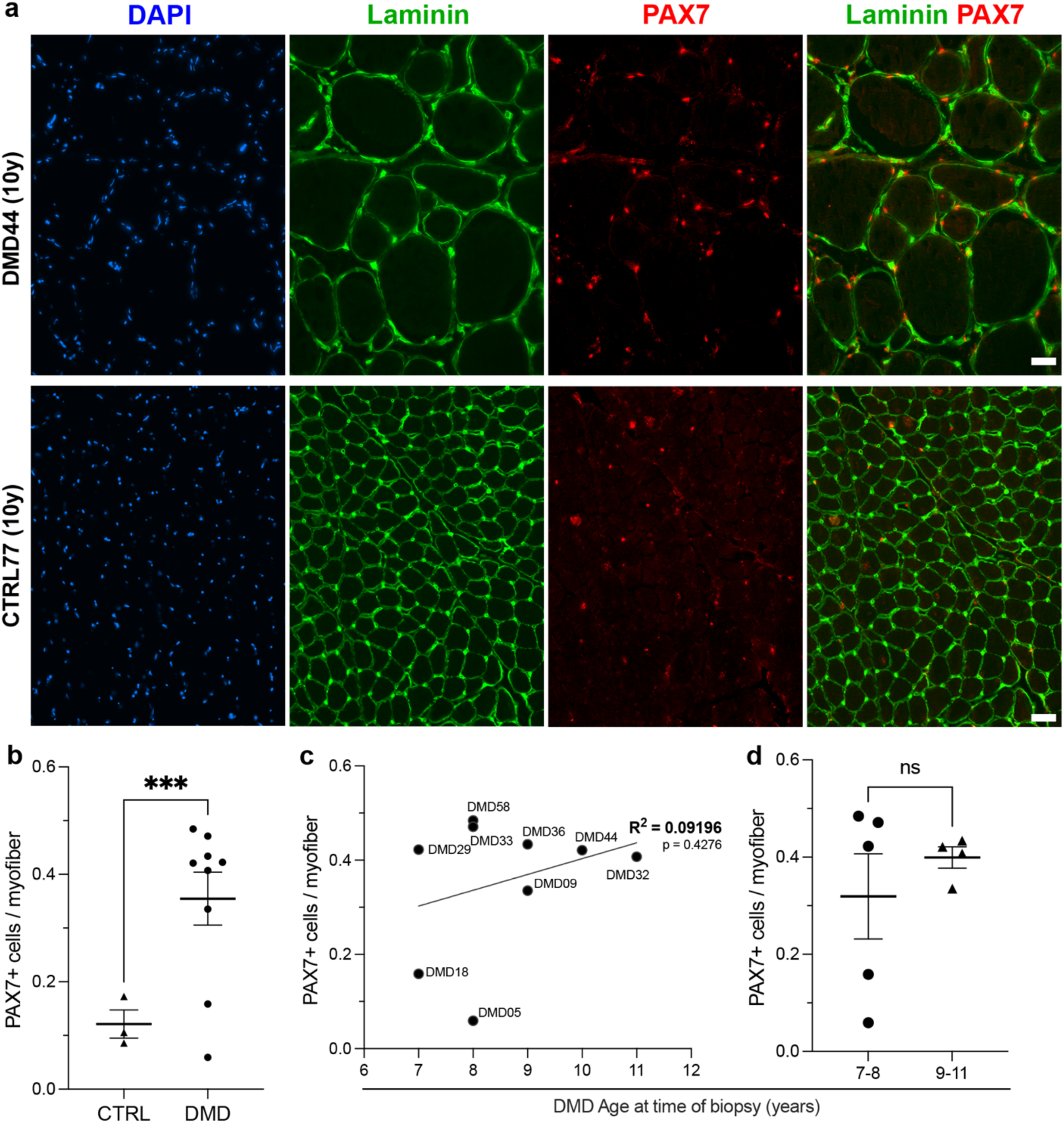
PAX7^+^ satellite cells are increased in DMD muscle. (**a**) Representative sections of DMD (DMD44, 10 years) and age-matched control (CTRL77, 10 years) muscle stained for DAPI (blue), laminin (green), PAX7 (red), and laminin/PAX7 overlay. (**b**) Quantification shows a significant increase in PAX7^+^ cells per myofiber in DMD compared with non-DMD controls (CTRL) (***P < 0.001, unpaired t test). (**c**) In DMD biopsies, the number of PAX7^+^ cells per myofiber shows a weak, non-significant correlation with age at time of biopsy (simple linear regression, R^2^ = 0.09196, p = 0.4276). (**d**) No significant difference in PAX7^+^ cells per myofiber is observed between younger (7–8 years) and older (9–11 years) DMD patients (ns, unpaired t test).

Taken together, using several standard approaches to quantify DMD disease progression, we observed little or no correlation with age likely due to the high variability between samples and patients.

### Determination of Regeneration Index

To normalize the variable degrees of myonecrosis and regeneration that occur within and between muscles in DMD patients, we developed and validated a measure of muscle regenerative capacity, which we term the Regenerative Index (RI). The RI is the ratio of newly formed (eMHC^+^) myofibers over necrotic (IgG^+^) myofibers.

To accurately quantify the dynamic balance between myonecrosis and regeneration we combined laminin immunostaining with staining for IgG or eMHC (Fig 4. a). Following the same laminin trace analysis performed to calculate CSA and MFF, regenerating myofibers were identified and enumerated based on eMHC expression and necrotic myofibers were identified and enumerated based on IgG staining. The complete list of quantified values and calculated RI are listed in Table 2.

**Figure 4.**
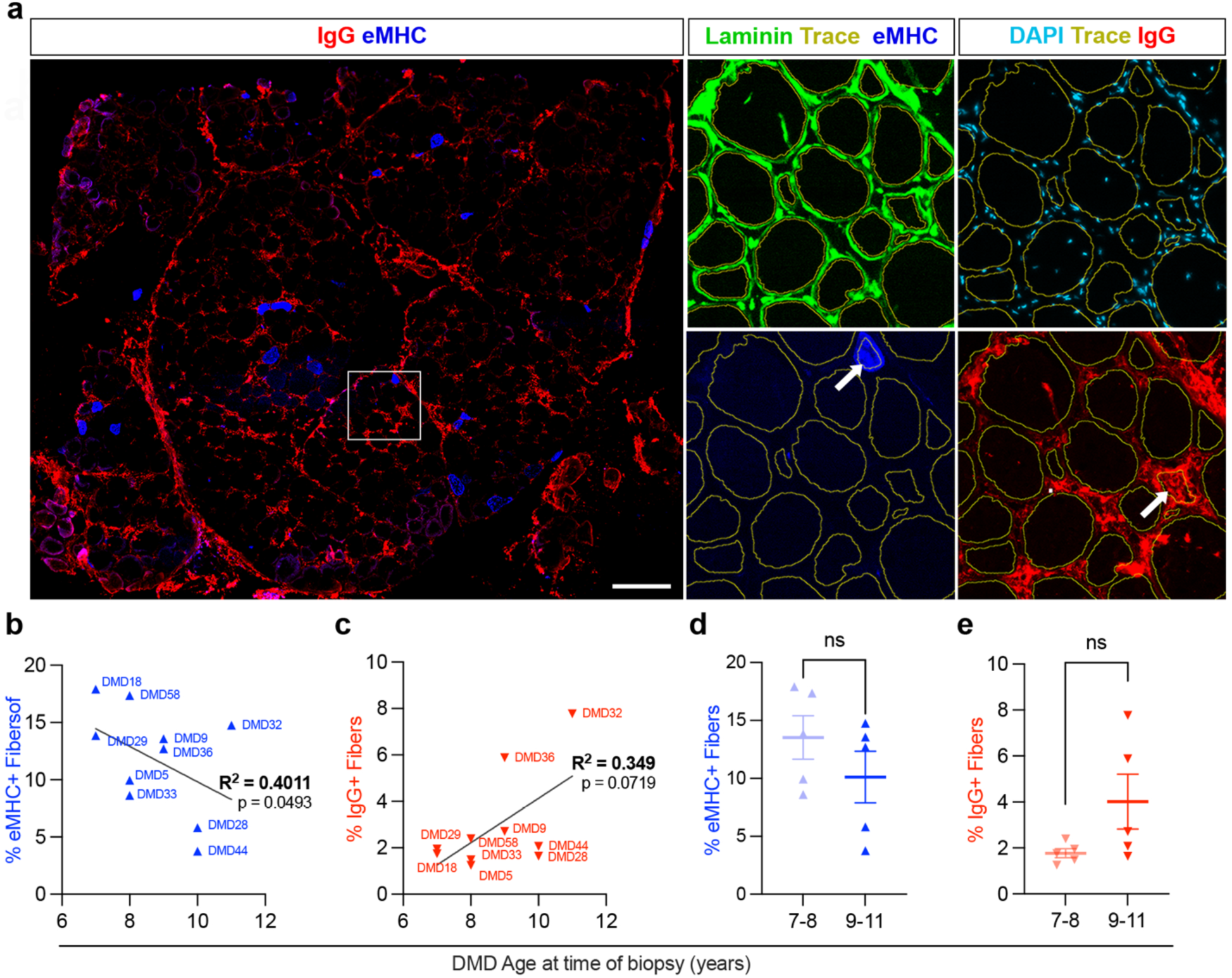
Decreased regeneration and increased degeneration in DMD muscle. (**a**) Representative immunofluorescence image of a DMD muscle biopsy stained for IgG (red) and embryonic myosin heavy chain (eMHC, blue), with 500 µm insets showing laminin (green) and automated tracing traces outlining individual fibers (yellow) to determine presence or absence of either eMHC or IgG (see thick white arrows). (**b**) Percentage of eMHC^+^ fibers in individual DMD patients plotted against age at biopsy, showing a weak but statistically significant positive correlation (R^2^ = 0.4011, p = 0.0493). (**c**) Percentage of IgG^+^ fibers in DMD patients plotted against age at biopsy, indicating a trend toward increased IgG^+^ fibers with age that does not reach statistical significance (R^2^ = 0.349, p = 0.0719). (**d, e**) Comparison of younger (7–8 years) and older (9–11 years) DMD patients shows no significant differences in the proportion of eMHC^+^ fibers (**d**) or IgG^+^ fibers (**e**) (ns, unpaired t test).

**Table 2.** Regenerative Index quantification for 7-11 year DMD samples.

| Sample | Age | IgG<sup>+</sup><br>fibers | eMHC<sup>+</sup><br>fibers | RI | Total<br>fibers | Percent<br>IgG<sup>+</sup><br>fibers | Percent<br>MHC<sup>+</sup><br>fibers |
| --- | --- | --- | --- | --- | --- | --- | --- |
| DMD 18 | 7 | 48 | 492 | 10.3 | 2747 | 1.75 | 17.91 |
| DMD 29 | 7 | 29 | 205.5 | 7.1 | 1484 | 1.95 | 13.85 |
| DMD 33a | 8 | 19 | 131 | 6.9 | 1520 | 1.25 | 8.62 |
| DMD 58a | 8 | 26 | 189 | 7.3 | 1089 | 2.39 | 17.36 |
| DMD 05b | 8 | 13 | 87 | 6.7 | 874 | 1.49 | 9.95 |
| DMD 09 | 9 | 21 | 105 | 5.0 | 774 | 2.71 | 13.57 |
| DMD 36 | 9 | 40.5 | 87.5 | 2.2 | 689 | 5.88 | 12.70 |
| DMD 28 | 10 | 38 | 135 | 3.6 | 2324 | 1.64 | 5.81 |
| DMD 44 | 10 | 17 | 31 | 1.8 | 822.5 | 2.07 | 3.77 |
| DMD 32 | 11 | 156 | 324 | 2.1 | 2195 | 7.11 | 14.76 |
| CTRL 12 | 8 | 0 | 0 | undefined | 6771 | 0.00 | 0.00 |
| CTRL 77a | 10 | 10 | 3 | 0.3 | 5721 | 0.17 | 0.05 |
| CTRL 76 | 11 | 5 | 0 | 0.0 | 4737 | 0.11 | 0.00 |

Enumeration of regenerating (eMHC^+^) or necrotic (IgG^+^) revealed age-related trends in muscle regeneration and degeneration. The percentage of eMHC^+^ myofibers showed a significant negative correlation with age at biopsy (R^2^ = 0.4011, p=0.0493), indicating moderate association between increasing age and a reduced proportion of regenerating myofibers (Figure 4b). In contrast, the percentage of IgG^+^ myofibers showed a weaker but positive correlation with age (R^2^ = 0.349, p = 0.0719) but did not reach statistical significance (Figure 4c). Comparisons between younger (7-8 years) and older (9-11 years) groups showed no statistically significant differences in either eMHC^+^ or IgG^+^ myofiber percentages (Figure 4d,e).

In striking comparison, determination of Regenerative Index (RI), defined as the ratio of newly forming eMHC^+^ myofibers to necrotic IgG^+^ myofibers (Figure 5a), revealed a strong inverse relationship with age at biopsy in boys with DMD (Figure 5b). The RI decreased significantly with age (R^2^ = 0.789, p = 0.0006), indicating a highly significant loss of regenerative capacity as boys grew older (Figure 5b). Furthermore, comparison between age groups revealed that 7–8-year-old boys had a markedly higher RI than 9–11-year-olds, with the difference reaching high statistical significance (Figure 5c).

**Figure 5.**
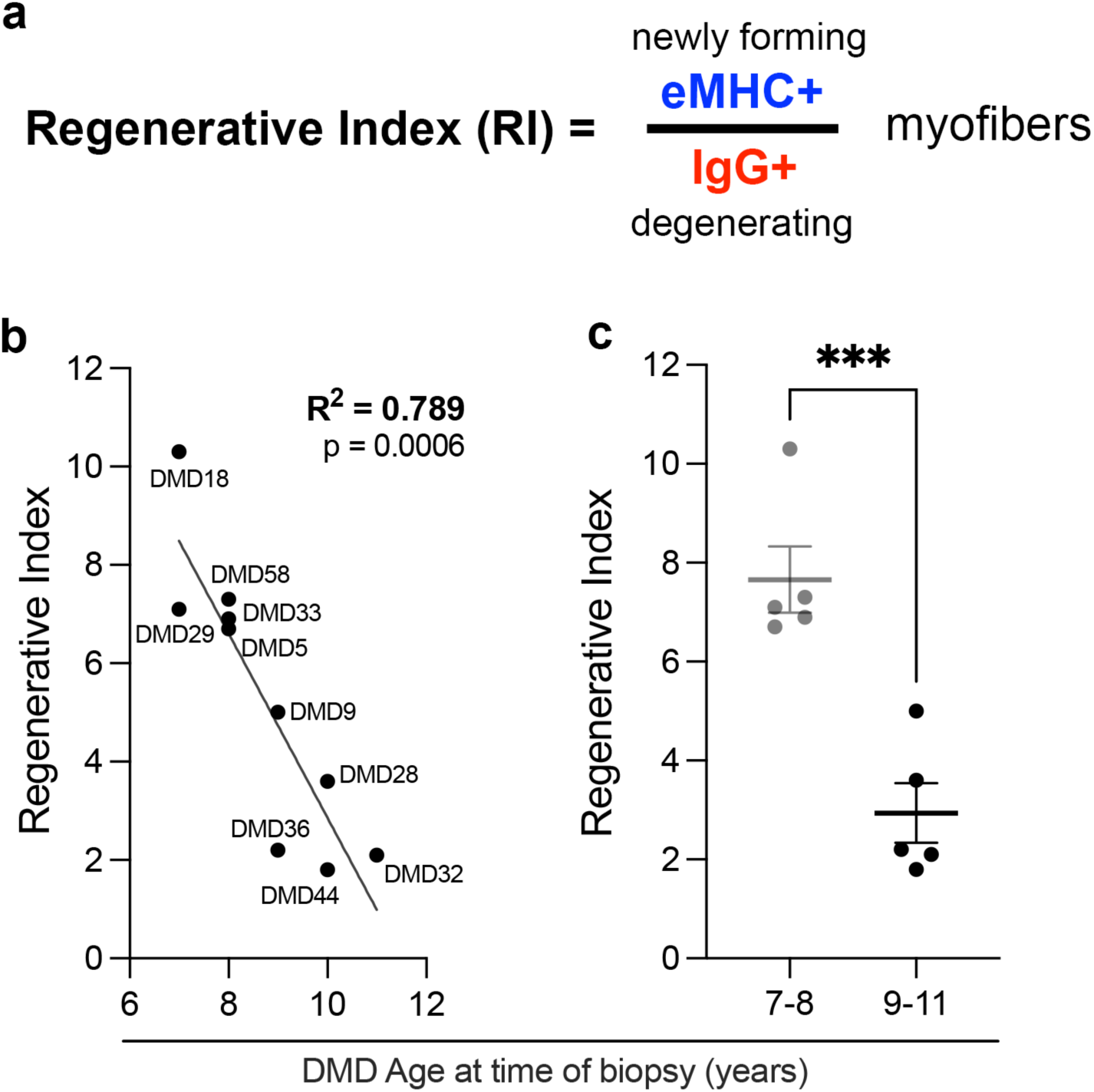
Regenerative Index is significantly decreased in DMD boys between the ages of 7 to 11 years. (**a**) Schematic representation of the Regenerative Index (RI), defined as the ratio of newly forming eMHC-positive myofibers to necrotic IgG-positive myofibers. (**b**) Plot of RI versus age at biopsy in DMD patients, showing a strong negative correlation between regenerative index and age (R^2^ = 0.789, p = 0.0006; simple linear regression). (**c**) Comparison of RI between younger (7–8 years) and older (9–11 years) DMD patients demonstrates a significantly higher regenerative index in the younger group (***p < 0.001, unpaired t test).

This finding demonstrates that the regenerative potential in DMD muscle declines sharply with age, as measured by the Regenerative Index. The RI being a ratio, is effective at normalizing the variation found between biopsies and thus provides a useful approach to evaluate the efficiency of regeneration. Therefore, we propose that determination of RI will allow the monitoring of changes in the ability of DMD muscle to regenerate, providing a powerful tool to assess treatments designed to enhance muscle regeneration.

## Discussion

The sharp decline in the Regenerative Index (RI) between 7 and 11 years of age, despite relatively stable myofiber morphology and satellite cell numbers, supports the potential of the RI as an effective measure of muscle regeneration. Moreover, the significant inverse correlation between RI and patient age suggests that the muscle’s ability to regenerate myofibers relative to ongoing myonecrosis deteriorates rapidly early in DMD. This early impairment in the efficiency of regenerative ability may correlate with the previously observed loss of satellite cell polarity in the absence of dystrophin [9, 17, 28]. These results underscore the need for treatments aimed at improving muscle regeneration in DMD.

In our studies, DMD satellite cell numbers per muscle fiber were increased relative to non-DMD controls without a significant reduction in abundance with increasing age. This aligns with earlier evidence suggesting that numbers of satellite cells remain elevated until much later in DMD [7]. However, the lack of dystrophin impairs satellite cell polarity resulting in their reduced ability to generate progenitors through asymmetric divisions[9]. A dysfunction in asymmetric satellite cell division may increase symmetric divisions, temporarily elevating total satellite cell numbers. However, if progenitor satellite cells are reduced in parallel, the expected long-term result is impaired muscle regeneration. A second negative effect of impaired satellite cell polarity may be increased chromosomal misalignment and mitotic catastrophe leading to satellite cells entering programmed cellular senescence [7, 9]. Correcting this polarity defect would maintain the number of progenitors associated muscle fiber regeneration. The RI measurement described here is a means by which therapeutics targeting DMD satellite cells may be evaluated.

One limitation of the Regenerative Index is the inability to compare DMD muscle samples directly to non-DMD, no diagnostic abnormality controls. As these controls typically display no eMHC^+^ or IgG^+^ myofibers, the RI could be undefined, zero, or based on a very small number of myofibers. Conversely, therapies targeting dystrophin expression can be directly evaluated against placebo controls. Restoration of dystrophin to levels observed in healthy muscle represents the ultimate therapeutic goal. Dystrophin can be measured in both by either Western blot or quantitative immunofluorescence to determine therapeutic success. However, unlike the dystrophin measurement, muscle regeneration markers cannot be compared between DMD and healthy individuals, due to the inherent differences in regeneration activity and pathology between diseased and normal muscle. Regenerative marker expression reflects ongoing disease processes rather than basal healthy muscle status. This distinction underscores the complexity of using RI as treatment benchmarks and shows the importance of longitudinal evaluation over time within DMD patients.

Therapies targeting satellite cell biology are emerging that may have potential in combination to therapies reintroducing dystrophin [29–31]. Evaluating such therapies will benefit from a method to more directly measure efficacy of increasing regenerative activity. Previously it was shown that increased myofiber size was observed comparing pre- and post-treatment muscle biopsies [26]. However, knowing myofiber size also increases over time without therapeutic intervention makes this difficult to interpret. Thus, morphometric analysis falls short of proving the increase is from drug treatment alone. As demonstrated, RI declines from age 7 to 11, therefore an increase after treatment or even a maintenance of RI would be a convincing measure of efficacy. Future longitudinal studies correlating RI with functional outcomes as well as examining RI across other muscle diseases and age groups will be important to validate and refine its use in clinical settings. Moreover, integrating transcriptomic and metabolic profiling with morphological indices like the Regenerative Index could refine our understanding of the mechanistic transitions underlying early regeneration failure in dystrophic muscle.

## Conclusions

The Regenerative Index (RI) integrates both regenerative and degenerative states to assess the net regenerative potential of muscle. Compared to methods currently in use, RI may provide a quantitative measure of muscle regeneration that accurately reflects ongoing regenerative activity in DMD. The age associated RI difference observed in our study reinforces the usefulness of the Index but also highlights an early decline in regenerative potential despite the sustained satellite cell pool. Therapeutics to improve this regenerative capacity in DMD are a much-needed addition to future treatments. The Regenerative Index provides a robust quantitative measure for evaluating the effectiveness of therapies aimed at improving muscle regeneration in conditions such as DMD.

## List of abbreviations

AAK1: Adaptor-associated protein kinase 1
BMD: Becker Muscular Dystrophy
CTRL: control (non-DMD)
CSA: cross-sectional area
DAPI: 4′,6-diamidino-2-phenylindole
DMD: Duchenne Muscular Dystrophy
DSHB: Developmental Studies Hybridoma Bank
eMHC: Embryonic Myosin Heavy Chain
H&E: Haematoxylin and Eosin
IgG: Immunoglobulin G
MFF: minimal fiber Feret (minimal Feret’s diameter)
Pax7/PAX7: paired box protein 7 (satellite cell marker)
PBS: phosphate-buffered saline
RI: Regenerative Index
RT: room temperature

## DECLARATIONS

### Ethics approval and consent to participate

All muscle samples used in the study are biopsies from the Repository in the University of Iowa Wellstone Muscular Dystrophy Specialized Research Center. Approval to use them for research is covered in the IRB protocol ID#200510769 (original protocol approval on 02/16/2006; most recent continuing review approval on 11/11/2025)

### Availability of data and materials

All data needed to evaluate the conclusions in the paper are present in the paper. Any additional information required to reanalyze the data reported in this work paper is available from the lead contact.

### Authors’ contributions

JKS and MAR designed research, analyzed data, and wrote the paper. CAM performed the RI and JGM the Pax7 staining and quantification. SSSR made contributions to the conception, data analysis and manuscript review. SAM provided all muscle biopsy cryosections, images of the histopathology, assisted with interpretation of the data, and edited the manuscript.

## Acknowledgments

The authors thank the University of Ottawa Imaging Core for use of microscopes and training required for scanning in entire muscle biopsy sections.

