## Supplementary Information for "Regenerative Index reveals declining muscle regeneration in paediatric patients with Duchenne muscular dystrophy"

Figures S1-S3

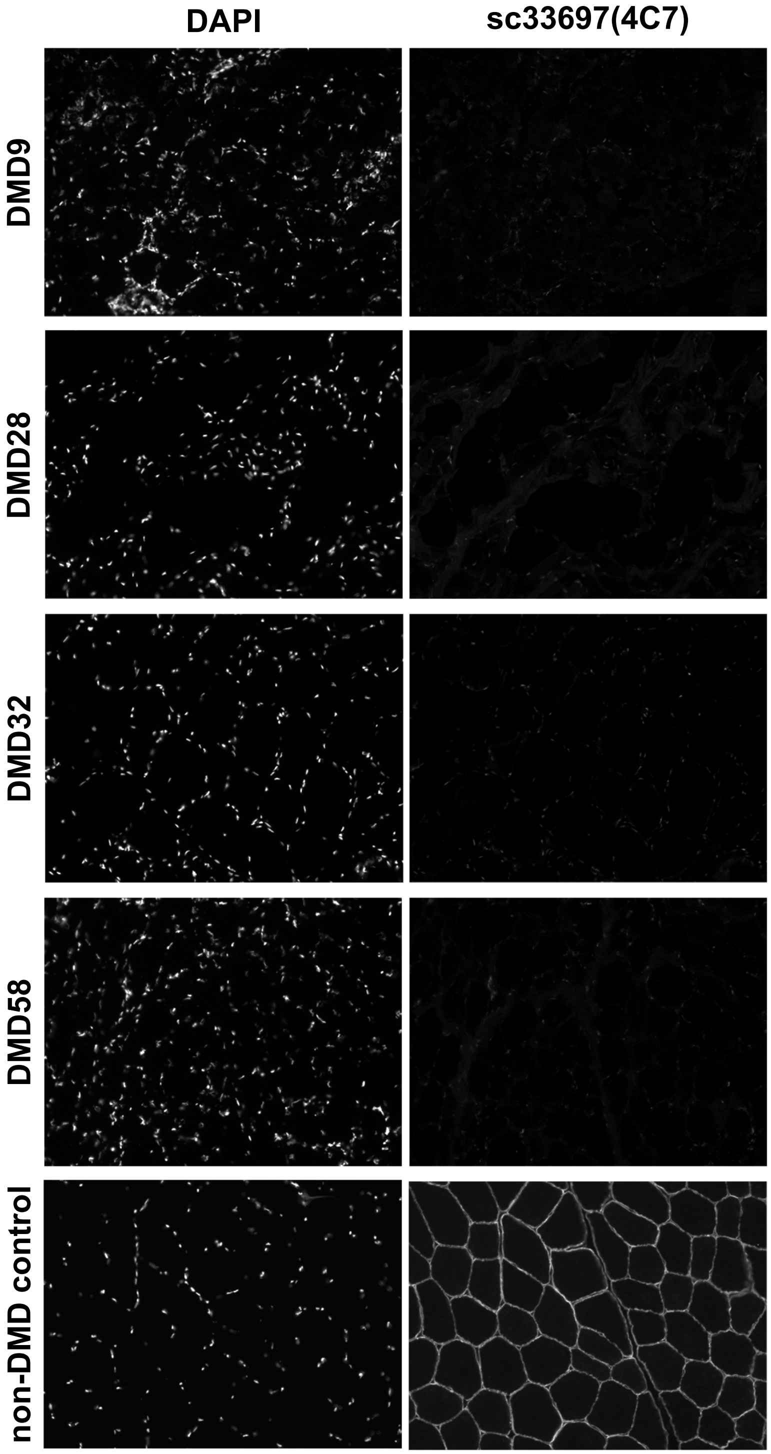


**Supplemental Figure 1. Representative images of dystrophin expression in a non-DMD and DMD patients of different ages.** Sections were stained with DAPI to visualize nuclei (left panels). Dystrophin immunostaining (right panel) shows a clear presence of dystrophin in the non-DMD control and absence in DMD samples. Scale bar represents 100 μm.

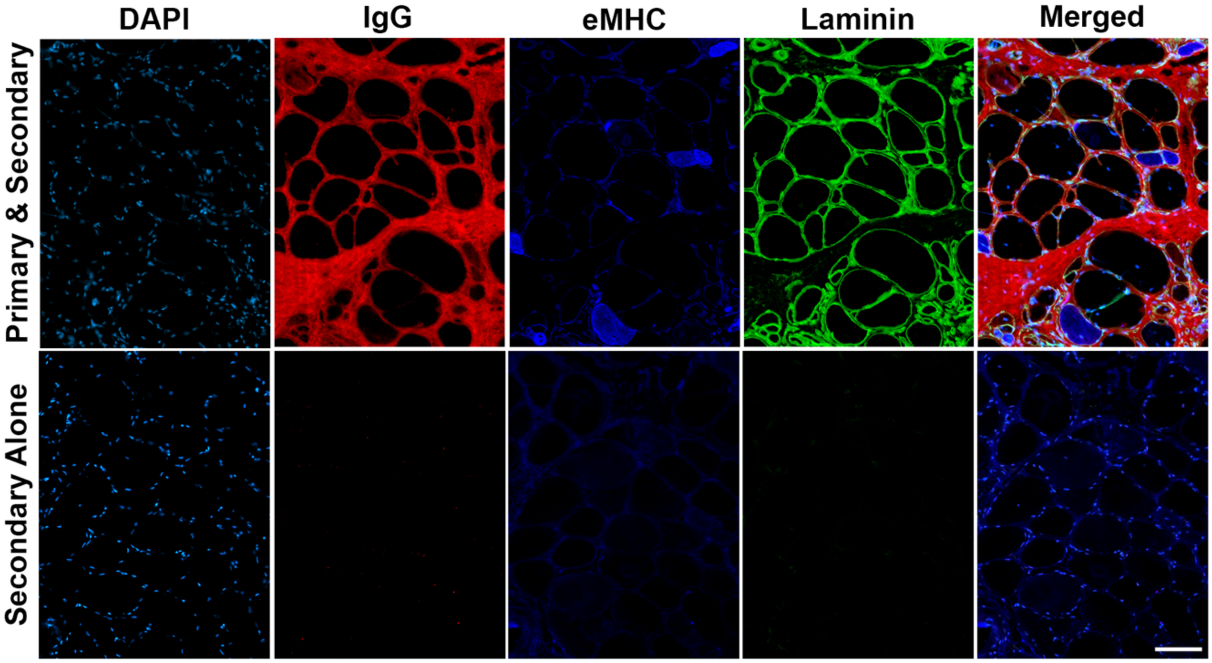


**Supplemental Figure 2. Representative immunofluorescence staining and secondary antibody–only controls in DMD32**. Sections were stained with DAPI (nuclei, cyan), anti-human IgG (red), anti-embryonic myosin heavy chain (eMHC, blue), and anti-laminin (green), and a merged image is shown in the right column for the full primary plus secondary antibody condition (top row). In the DAPI and secondary antibody only condition (bottom row), no specific signal is detected, demonstrating minimal background and confirming the specificity of the primary antibody staining. Scale bar represents 100 μm.

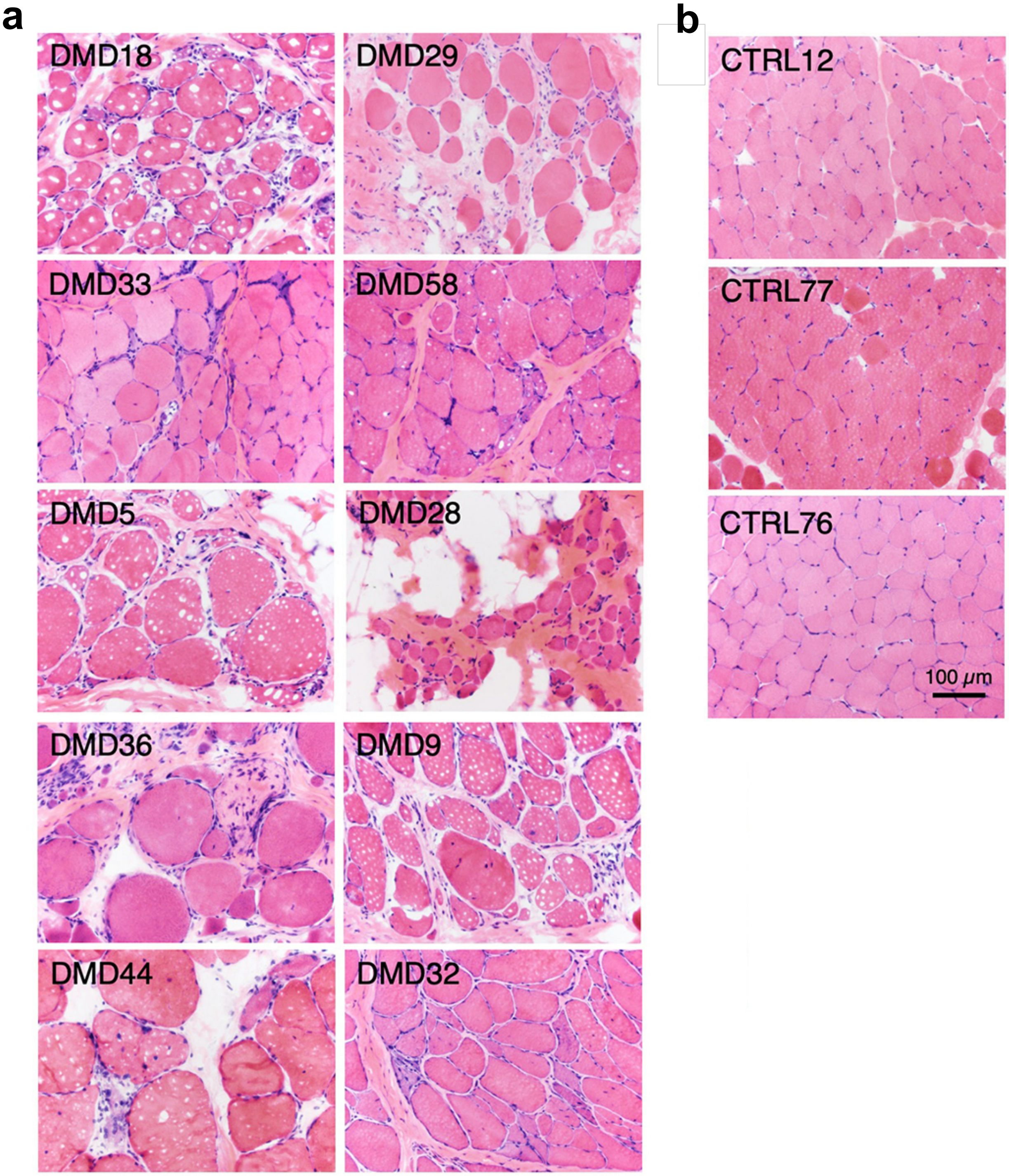


**Supplemental Figure 3. Representative H&E photomicrographs from 13 muscle biopsy sections used in the study.** (**a**)10 DMD muscle biopsy sections and (**b**) 3 non-DMD controls. Scale bar represents 100 μm.
